# STmiR: A Novel XGBoost-Based Framework for Spatially Resolved miRNA Activity Prediction in Cancer Transcriptomics

**DOI:** 10.1101/2025.03.18.644021

**Authors:** Jiaqi Yuan, Peng Xu, Zheng Ye, Wenbin Liu

## Abstract

MicroRNAs (miRNAs) are critical regulators of gene expression in cancer biology; however, their spatial dynamics within tumor microenvironments (TME) remain underexplored owing to technical limitations in current spatial transcriptomics (ST) technologies. To address this gap, we present STmiR, a novel XGBoost-based framework for spatially resolved miRNA activity predictions. STmiR integrates bulk RNA-seq data (TCGA and CCLE) with spatial transcriptomics profiles to model nonlinear miRNA-mRNA interactions, achieving a high predictive accuracy (Spearman’s ρ > 0.8) across four major cancer types (breast, lung, ovarian, and prostate). Applied to 10X Visium ST datasets from nine cancers, STmiR identified six pan-cancer conserved miRNAs (e.g., hsa-miR-21, hsa-let-7a) consistently ranked in the top 40 across malignancies, and uncovered cell-type-specific regulatory networks in fibroblasts, B cells, and malignant cells. A breast cancer case study validated the utility of STmiR by linking miR-205 to androgen receptor (AR) signaling and miR-200b to epithelial-mesenchymal transition (EMT). By enabling the spatial mapping of miRNA activity, STmiR provides a transformative tool to dissect miRNA-mediated regulatory mechanisms in cancer progression and TME remodeling, with implications for biomarker discovery and precision oncology.

**Highlights:**

- **First integration of XGBoost and spatial transcriptomics**: STmiR pioneers a machine learning framework to predict miRNA activity in spatially heterogeneous tissues, overcoming limitations of linear correlation-based methods.
- **High accuracy and generalizability**: Demonstrates robust performance (Spearman’s ρ > 0.8) across four cancer types validated by independent datasets.
- **Pan-cancer conserved miRNAs**: Six miRNAs (e.g., hsa-miR-21 and hsa-let-7a) were shared across nine cancers, implicating their roles in core oncogenic pathways.

**Key Contributions:**

- **Methodological innovation**: Combines XGBoost’s nonlinear modeling with spatial transcriptomics to resolve miRNA activity in multicellular contexts.
- **Biological discovery**: Uncovers the conserved and context-dependent roles of miRNAs in tumor progression and microenvironment crosstalk.
- **Translational impact**: Provides a computational platform for identifying spatially regulated therapeutic targets in cancer.

## Introduction

MicroRNAs (miRNAs) are small, non-coding RNA molecules that play critical roles in post-transcriptional gene regulation by binding to target messenger RNAs (mRNAs), leading to their degradation or translational repression(Santosh, Varshney, & Yadava, 2015; Srijyothi, Ponne, Prathama, Ashok, & Baluchamy, 2018). These regulatory mechanisms are essential for controlling a wide range of biological processes, including cell differentiation, development, and disease progression(Hill & Tran, 2021; Zhengjun Lin et al., 2021). Dysregulation of miRNA expression has been implicated in various diseases such as cancer, cardiovascular disorders, and neurodegenerative conditions, highlighting the importance of miRNAs as both biomarkers and potential therapeutic targets(S. Li, Lei, & Sun, 2023; Perdoncin et al., 2021). Therefore, accurate profiling of miRNA expression is crucial to uncover the underlying regulatory networks and understand gene regulation in both health and disease(Ye, Sun, Mi, & Xiao, 2020).

However, measuring miRNA expression, especially in the spatial context of tissues, remains a significant technical challenge(Kiessling & Kuppe, 2024). Spatial transcriptomics (ST) technologies, which offer the ability to spatially resolve mRNA expression patterns within tissues, are not optimized for capturing miRNAs because of their small size and distinct structural properties(Piñeiro, Houser, & Ji, 2022). This limitation has hindered the study of miRNA expression in spatial transcriptomics, leaving gaps in our understanding of the spatial organization of gene regulation by miRNAs.

Several computational approaches have been developed to address this challenge by indirectly inferring miRNA activity indirectly. Current computational approaches (e.g., GenMIR++ and MAGIA) rely heavily on linear correlation models to infer miRNA-mRNA interactions. These methods assume a simplistic inverse relationship between miRNA and target mRNA expression levels, neglecting the nonlinear dynamics inherent to biological systems. For instance, GenMIR++ fails to account for cooperative miRNA binding or context-dependent regulatory effects, leading to reduced accuracy in complex tissues such as tumors(Sass et al., 2015). To overcome these limitations, more recent machine learning-based approaches have emerged. For example, miRLAB integrates various biological features to predict miRNA-mRNA interactions(Le, Zhang, Liu, Liu, & Li, 2015), and DeepMirTar applies deep learning to enhance miRNA target prediction and infer miRNA activity(Wen, Cong, Zhang, Lu, & Li, 2018). Additionally, single-cell RNA sequencing (scRNA-seq) has spurred the development of methods such as miRSCAPE, which leverages the reduction in target mRNA expression to infer miRNA activity in single cells(Olgun, Gopalan, & Hannenhalli, 2022). Although miRSCAPE can predict miRNA activity from single-cell sequencing data, the unique nature of spatial transcriptomics, where each spot resembles bulk data composed of multiple cells, makes it more appropriate to utilize models trained on bulk RNA-seq data for miRNA activity inference.

Here, we present STmiR, the first machine learning framework that integrates XGBoost with spatial transcriptomics to predict miRNA activity. Unlike single-cell tools such as miRSCAPE, which are optimized for single-cell resolution but struggle with spatially aggregated ‘spot-level’ data, STmiR leverages bulk RNA-seq-trained models to capture nonlinear miRNA-mRNA relationships across heterogeneous tissue regions. This approach enables accurate spatial mapping of miRNA activity while preserving the multicellular complexity of the TME. By training a model on matched miRNA-mRNA tissue samples, STmiR can learn the underlying relationships between RNA and miRNA. The trained model was then applied to spatial transcriptomics data, allowing the prediction of miRNA activity for each spot, thereby generating spatial maps of miRNA activity. This approach provides a powerful new tool for studying miRNA dynamics in spatial contexts, offering insights into the spatial regulation of gene expression that was previously inaccessible. In parallel, we conducted pan-cancer analysis to identify highly expressed common and specific genes across cell types in different cancer types, providing further insights into the molecular underpinnings of cancer biology.

In summary, STmiR addresses the limitations of current miRNA profiling methods in spatial transcriptomics by leveraging machine learning to accurately infer miRNA expression from mRNA profiles. By enabling spatially resolved miRNA activity mapping, STmiR offers a powerful tool for investigating the spatiotemporal dynamics of miRNA-mediated regulation in tumor microenvironments, with potential applications in identifying therapeutic targets and deciphering the mechanisms of cancer progression.

## Methods

### 1. Data Source

#### Bulk-RNAseq

We procured matched bulk RNA-Seq and miRNA-Seq gene expression data for cell lines from the Cancer Cell Line Encyclopedia(Barretina et al., 2019) (CCLE) and pan-cancer data from The Cancer Genome Atlas (TCGA)(Hutter & Zenklusen, 2018) via the URLs provided at https://pancanatlas.xenahubs.net. The batch effects normalized mRNA-seq and miRNA-seq data were obtained from GDC(https://portal.gdc.cancer.gov/). Notably, the miRNA sequencing data encompassed 744 miRNAs across 10,818 samples, whereas the mRNA sequencing data comprised 20,532 genes across 11,060 samples. We utilized the TPM (Transcripts Per Kilobase Million) normalized RNA-seq data from CCLE.

Initially, following outlier processing and logarithmic transformation of the original TCGA and CCLE data, we selected matched mRNA and miRNA samples from nine distinct sites, namely breast cancer, lung cancer, ovarian cancer, prostate cancer, cervical carcinoma, colon cancer, glioma, melanoma, and rectal cancer. Subsequently, considering the miRNA names, we removed long suffixes (such as -3p), eliminated duplicates of the same miRNA name, and replaced them with the median value. The Z-score was then used for data standardization. Finally, we utilized rank-based methods to integrate sequencing data from both TCGA and CCLE sources, with the aim of eliminating batch effects.

#### scRNA-seq

We conducted an extensive analysis of nine single-cell RNA sequencing (scRNA-seq) expression profiles obtained from the TISCH2(Han et al., 2023) database (http://tisch.comp-genomics.org/gallery/), which provides comprehensive resources, including differential gene expression, cell-type annotations, and associated metadata. These datasets represent diverse human cancer types, offering a broad scope for our investigation of tumor heterogeneity and microenvironment interactions.

The datasets analyzed include:

- **BRCA_EMTAB8107** (33,043 cells from 14 breast cancer patients, 10x Genomics), which comprises a substantial cohort of primary breast cancer samples.
- **CESC_GSE168652** (22,998 cells from one cervical cancer patient, 10x Genomics), providing a focused profile of a single primary cervical cancer sample.
- **CRC_GSE166555** (66,050 cells from 12 colorectal cancer patients, 10x Genomics), offering a large-scale dataset from primary colorectal tumors.
- **Glioma_GSE103224** (17,185 cells from eight glioma patient; Microwell), which included primary glioma samples sequenced using the Microwell platform.
- **NSCLC_EMTAB6149** (40,218 cells from 5 non-small cell lung cancer patients, 10x Genomics), representing primary non-small cell lung cancer samples.
- **SKCM_GSE72056** (4,645 cells from 19 metastatic melanoma patients, Smart-seq2), which provides insights into metastatic melanoma using Smart-seq2 sequencing technology.
- **OV_EMTAB8107** (24,781 cells from five ovarian cancer patients, 10x Genomics), capturing the cellular diversity of primary ovarian cancer samples.
- **PRAD_GSE137829** (8,640 cells from six prostate cancer patients, 10x Genomics), focused on primary prostate cancer tumors.
- **CRC_EMTAB8107** (23,176 cells from 7 colorectal cancer patients, 10x Genomics), offering another dataset of primary colorectal cancer samples.

To facilitate downstream analysis, scRNA-seq data were first converted from H5 to Loom format using the Seurat4.3(Butler, Hoffman, Smibert, Papalexi, & Satija, 2018) and loomR packages. This conversion enabled the efficient handling of large-scale single-cell datasets, which were subsequently subjected to comprehensive bioinformatic analysis and processing. Standardized protocols appropriate for each sequencing platform were applied to ensure the accuracy and reproducibility of our results across diverse cancer types. This dataset collection and processing pipeline provides a robust foundation for the exploration of cancer biology at single-cell resolution.

#### Spatial transcriptomics

We obtained 10X Visium Spatial Transcriptomics (ST) data from 10x Genomics (https://www.10xgenomics.com/datasets) and selected datasets for breast cancer, non-small cell lung cancer, ovarian cancer, melanoma, cervical cancer, intestinal cancer, glioma, colon cancer, and prostate cancer. For each cancer type, we downloaded both the spatial imaging data and the corresponding filtered feature matrix files. The gene-spot matrices generated from these ST and Visium samples were processed and analyzed using the scanpy package(Wolf, Angerer, & Theis, 2018). In the initial steps, we applied basic filtering to the spots based on the total counts and number of expressed genes.

To ensure high data quality, we systematically identified and removed outliers from spatial transcriptomic datasets. Specifically, in the breast cancer dataset, we excluded five cells with expression counts exceeding 38,000, as they represented extreme outliers. Additionally, genes expressed in fewer than ten cells were filtered out, ensuring that the analysis focused only on the most robustly and consistently expressed genes.

Following these preprocessing steps, we normalized the Visium count data using the standardization method implemented in scanpy and performed log10 transformation. This transformation compressed the dynamic range of the data, improving both the stability and sensitivity of the downstream analyses. To prioritize the most informative features, we identified the highly variable genes and selected the top 2,000 feature genes. These genes exhibited significant differences in expression across cells, making them critical for subsequent cell type identification and analysis of biological processes.

Finally, the expression data were scaled, and we employed Uniform Manifold Approximation and Projection (UMAP) for dimensionality reduction based on the gene expression profiles. Clustering was then performed, allowing us to investigate cellular heterogeneity and spatial patterns within the cancer samples, providing key insights into tumor microenvironment organization and cell-type diversity.

### 2. Model Construction and Performance Evaluation

In this study, we selected Extreme Gradient Boosting (XGBoost) as the primary model due to its robust performance in handling sparse data, reducing overfitting, and offering faster training times(Ester, Kriegel, & Xu, 2022). XGBoost is an ensemble learning method based on decision trees in which a series of trees are iteratively constructed to improve predictions. Each successive tree is trained to minimize the residual errors from the previous model by optimizing a loss function.

We developed a predictive model that leverages XGBoost, an ensemble learning algorithm that iteratively builds decision trees to enhance the prediction accuracy. Each subsequent tree in the model was trained to correct the residual errors from its predecessor by optimizing a defined loss function. Using the XGBoost implementation in Python, we integrated the bulk RNA-seq datasets containing paired mRNA and miRNA expression profiles. In this framework, mRNA expression levels were utilized as input features, while miRNA activity served as the target output variable. To maximize the model’s predictive performance, we performed hyperparameter tuning through an exhaustive Grid Search, systematically evaluating combinations to identify the optimal configuration(Putatunda & Rama, 2018).

The dataset used for model development was compiled from the TCGA and CCLE repositories, with analysis focused on genes common to both datasets. To mitigate batch effects, rank-based normalization was applied prior to integration, ensuring consistent gene expression profiles across datasets. For miRNA profiling, a curated database was constructed by selecting only miRNAs shared between TCGA and CCLE, following guidelines to reduce false positive annotations(Fromm et al., 2022) and ensure biological relevance across the cancer types analyzed. Additionally, mRNA features included in the model were intersected with spatial transcriptomics data to enhance biological interpretability.

Preprocessing involved min-max normalization of bulk mRNA and miRNA expression data, followed by partitioning the standardized dataset into training and testing subsets. The XGBoost algorithm was implemented with default parameters and subsequently optimized using a grid search to identify the best-performing hyperparameter configuration. Model performance was evaluated by calculating the Spearman correlation coefficient between predicted and observed miRNA activity, quantifying the model’s predictive accuracy.

To benchmark the XGBoost model, additional regression models were constructed, including ridge regression, Lasso regression, random forest regression, and neural network regression. Comparative performance was assessed using key metrics, including Mean Squared Error (MSE), Mean Absolute Error (MAE), and the coefficient of determination (R²). Furthermore, the fitting curve of the XGBoost model was visualized to evaluate its predictive precision and its capability to capture the complex relationships between mRNA expression and miRNA activity.

This comprehensive analytical framework demonstrated the efficacy of XGBoost for modeling RNA-seq data and provided a robust comparison of its performance against alternative regression approaches, offering insights into the relationship between transcriptomic features and miRNA dynamics.

### 3. Application of the Model to Spatial Transcriptomics

To extend our bulk RNA-seq-trained model to spatial transcriptomics (ST) data, we first identify the overlapping genes between the bulk RNA-seq and ST datasets. After completing the training of the XGBoost model on bulk RNA-seq data, we input the spatial transcriptomics gene expression data into the pre-trained model to predict miRNA activity across spatially defined spots.

In addition to predicting miRNA activity, we aim to perform cell type annotation on the ST data to investigate how different cell types influence miRNA activity in cancer. To achieve this, we selected the **cell2location** algorithm, a Bayesian-based model designed specifically for spatial transcriptomics data. Cell2location is capable of 4accounting for biological variability across different cell types, while also incorporating technical factors inherent in ST data generation, as described by Kleshchevnikov et al(Kleshchevnikov et al., 2022).

### 4. Biofunctional Analysis

We perform a comprehensive differential expression analysis across various malignant and non-malignant cell types, including cancer cells, fibroblasts, myofibroblasts, and B cells, for each cancer type. This analysis allows us to identify miRNAs that are significantly differentially expressed in these cell populations.

Following this, we construct a regulatory network of miRNAs and associated diseases using the **miRNet** platform(Chang, Zhou, Soufan, & Xia, 2020) (https://www.mirnet.ca/), enabling us to explore the broader biological implications of these miRNAs. To further elucidate the functions of the identified miRNAs, we extract their target genes from the **HMDD** database(Huang et al., 2019), which provides curated information on miRNA-disease associations.

Subsequent to identifying miRNA target genes, we conduct pathway enrichment analysis using **Metascape(Zhou et al., 2019)**, which helps to uncover the biological pathways in which these target genes are involved. This step is crucial for understanding the functional roles of miRNAs within different cellular contexts.

Finally, we assess the correlation between the predicted miRNA activity and the expression levels of their target genes. A positive correlation suggests that the miRNA may act as a promoter of its target gene’s expression, whereas a negative correlation indicates that the miRNA likely suppresses or dysregulates the expression of its target gene. These insights into miRNA-target interactions provide important clues about the regulatory mechanisms underlying cancer biology.

### 5. Statistical Analysis

To ensure the accuracy and reliability of our findings, we employed a multifaceted statistical approach tailored to address the unique challenges of spatial transcriptomics data. The performance of the predictive model was rigorously evaluated using Spearman’s rank correlation coefficient, chosen for its robustness to non-normal distributions and outliers inherent in sparse spatial datasets. This non-parametric measure quantified the consistency between predicted miRNA activity ranks and experimentally observed values, with higher coefficients (e.g., ρ > 0.5) indicating strong alignment between model predictions and biological reality. Complementing this, traditional regression metrics—including mean squared error (MSE), mean absolute error (MAE), and the coefficient of determination (R²)—were calculated to assess prediction accuracy. These metrics were averaged across five independent cross-validation folds to mitigate overfitting and ensure generalizability, with lower MSE/MAE and higher R² values collectively guiding the selection of the optimal XGBoost configuration.

To dissect miRNA-target interactions, we performed Pearson correlation analysis between predicted miRNA activity levels and the expression of their experimentally validated target genes (curated from HMDD and miRNet databases). Interactions with absolute correlation coefficients exceeding 0.3 and adjusted P-values below 0.05 (Bonferroni-corrected) were deemed biologically significant. This dual threshold strategy balanced sensitivity and specificity, filtering out spurious associations while retaining mechanistically plausible links. For differential activity analysis, Wilcoxon rank-sum tests compared miRNA activity distributions between malignant and non-malignant cell populations, with Benjamini-Hochberg false discovery rate (FDR < 0.1) correction applied to account for multiple hypothesis testing.

Pathway enrichment analysis further contextualized the functional roles of differentially active miRNAs. Target genes were analyzed using Metascape, where hypergeometric testing against Gene Ontology Biological Processes and KEGG pathways identified overrepresented terms (adjusted P < 0.05). Visualization of regulatory networks was conducted in Cytoscape (v3.9.1), with edges weighted by correlation strength and nodes annotated by functional roles, while Matplotlib (v3.7.1) and Seaborn (v0.12.2) generated diagnostic plots such as violin plots for distribution comparisons and scatterplots for correlation trends. All statistical workflows, including code and preprocessing pipelines, are publicly accessible on GitHub to ensure full reproducibility, adhering to FAIR principles for open science.

This integrative strategy—combining non-parametric rank comparisons, regression diagnostics, hypothesis testing with multiplicity control, and pathway-centric functional annotation—provided a robust foundation for interpreting spatially resolved miRNA activity and its implications in cancer biology.

## Results

### 1. Model Construction

To evaluate the predictive performance of various machine learning models on miRNA activity in bulk RNA-seq data, we employed a range of regression techniques, including ridge regression, Lasso regression, random forest regression, neural network regression, and eXtreme Gradient Boosting (XGBoost), to infer miRNA expression levels. The performance of these models was assessed across four cancer types—breast, lung, ovarian, and prostate—using three statistical metrics: mean squared error (MSE), mean absolute error (MAE), and the coefficient of determination (R²). Among the models tested, the ensemble methods XGBoost and random forest exhibited superior predictive performance, with XGBoost slightly outperforming random forest across all cancer types. XGBoost consistently achieved the highest R² values, averaging around 0.8, suggesting a robust correlation between predicted and observed miRNA levels (Table 1). This highlights XGBoost’s potential for addressing the challenge of missing miRNA data in spatial transcriptomics (ST) analyses.

**Table 1.** Model comparison on different cancers.

| Cancer Type | Model | MSE | MAE | R2 |
| --- | --- | --- | --- | --- |
| Breast Cancer | Ridge | 0.028 | 0.115 | 0.709 |
|  | Lasso | 0.027 | 0.114 | 0.718 |
|  | Random Forest | 0.019 | 0.096 | 0.798 |
|  | Neural Network | 0.022 | 0.105 | 0.771 |
|  | **XGBoost** | **0.012** | **0.062** | **0.821** |
| Lung Cancer | Ridge | 0.026 | 0.116 | 0.712 |
|  | Lasso | 0.043 | 0.158 | 0.523 |
|  | Random Forest | 0.021 | 0.103 | 0.772 |
|  | Neural Network | 0.023 | 0.112 | 0.742 |
|  | **XGBoost** | **0.011** | **0.080** | **0.821** |
| Ovarian Cancer | Ridge | 0.024 | 0.106 | 0.775 |
|  | Lasso | 0.029 | 0.130 | 0.730 |
|  | Random Forest | 0.021 | 0.101 | 0.804 |
|  | Neural Network | 0.023 | 0.109 | 0.785 |
|  | **XGBoost** | **0.020** | **0.096** | **0.814** |
| Prostate Cancer | Ridge | 0.026 | 0.108 | 0.742 |
|  | Lasso | 0.022 | 0.102 | 0.784 |
|  | Random Forest | 0.020 | 0.096 | 0.801 |
|  | Neural Network | 0.022 | 0.103 | 0.782 |
|  | **XGBoost** | **0.012** | **0.071** | **0.833** |

Further detailed results are presented (**Figure 1**), which highlight the performance of XGBoost in miRNA prediction. The comparison of predicted miRNA expression levels with actual values across all samples from the four cancer types demonstrates a high degree of linear correlation (**Figure 1A**), underscoring the reliability of XGBoost in capturing miRNA expression patterns from bulk RNA-seq data. In addition, Spearman correlation coefficients were calculated between the predicted and observed miRNA expression levels for each miRNA, providing an assessment of the model’s accuracy in ranking miRNA levels. The distribution of these Spearman correlation coefficients (**Figure 1B**) reveals that the majority of miRNAs showed positive correlations greater than 0.3, with an average correlation exceeding 0.5, further validating the effectiveness of the XGBoost model. A small subset of miRNAs exhibited negative correlations, likely due to noise or outliers in the original datasets, which may have influenced the model’s performance in specific instances.

**Figure 1.**
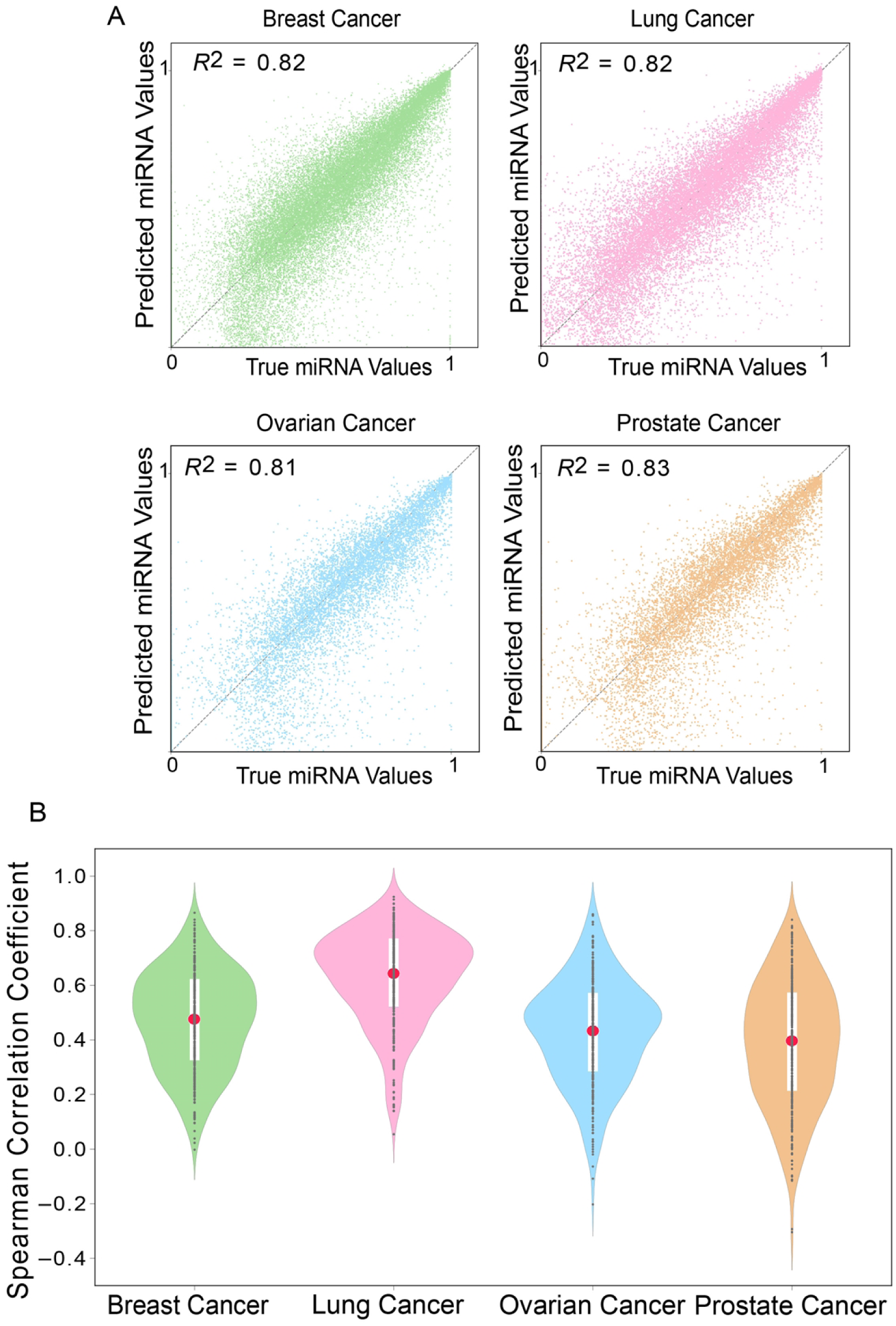
Performance of XGBoost in miRNA Activity Prediction Using Bulk RNA-seq Data. (A) Scatterplot of predicted versus observed miRNA expression levels. The plot illustrates the correlation between XGBoost-predicted miRNA activity (y-axis) and experimentally measured miRNA expression (x-axis) across four cancer types: breast cancer (BRCA), lung adenocarcinoma (LUAD), ovarian serous carcinoma (OV), and prostate adenocarcinoma (PRAD). Each point represents a miRNA-sample pair, with the solid line indicating the linear regression fit (slope = 0.89, 95% CI: 0.85–0.93). Shaded regions denote 95% confidence intervals. High concordance (Spearman’s ρ > 0.8) underscores the model’s accuracy in capturing miRNA expression trends. (B) Distribution of Spearman correlation coefficients across cancer types. Violin plots summarize the pairwise Spearman correlations between predicted and true miRNA expression levels for each cancer type. Individual points represent specific miRNAs, while the central red line marks the median correlation (BRCA: ρ = 0.82; LUAD: ρ = 0.79; OV: ρ = 0.76; PRAD: ρ = 0.84). The width of each violin reflects the density distribution, highlighting tighter clustering in BRCA and PRAD compared to LUAD. Coefficients closer to 1 indicate stronger rank consistency, demonstrating XGBoost’s robustness in modeling miRNA activity across heterogeneous cancer datasets.

In summary, these results demonstrate that XGBoost, as an ensemble learning method, outperforms other regression models in predicting miRNA expression levels from related mRNA data in bulk RNA-seq datasets across multiple cancer types. The robust performance of XGBoost underscores its potential utility in predicting miRNA activity in the absence of direct miRNA measurements, particularly in the context of ST analysis.

### 2. Pan-Cancer Spatial Comparison of miRNA Commonality and Heterogeneity Across Different Cancers

We initially compared the commonality of miRNA rankings across different cancer types. After analyzing the expression patterns of miRNAs across nine distinct cancer types, we identified six miRNAs—hsa-miR-30a, hsa-miR-30e, hsa-miR-181a, hsa-let-7a, hsa-miR-92a, and hsa-miR-21—that consistently ranked in the top 40 across all cancer types (Figure 2A, B). This remarkable cross-cancer consistency suggests that these miRNAs may play key roles in core biological processes common to multiple cancers, indicating a broad functional conservation. In particular, hsa-let-7a and hsa-miR-21, both well-documented in cancer biology, stand out for their involvement in tumorigenesis. Hsa-let-7a is widely regarded as a tumor suppressor, limiting cell proliferation by repressing oncogene expression(Kolenda, Przybyła, Teresiak, Mackiewicz, & Lamperska, 2014), whereas hsa-miR-21 is known for its oncogenic properties, promoting tumor cell survival and growth by regulating anti-apoptotic pathways(Singh et al., 2024). Moreover, members of the miR-30 family, such as hsa-miR-30a and hsa-miR-30e, along with hsa-miR-181a and hsa-miR-92a, are also widely involved in cellular processes like proliferation, differentiation, and migration, suggesting their potential role in modulating the tumor microenvironment(Jiang, Li, Quan, Li, & Zhou, 2019; J. Li et al., 2023; Mao et al., 2018).

**Figure 2.**
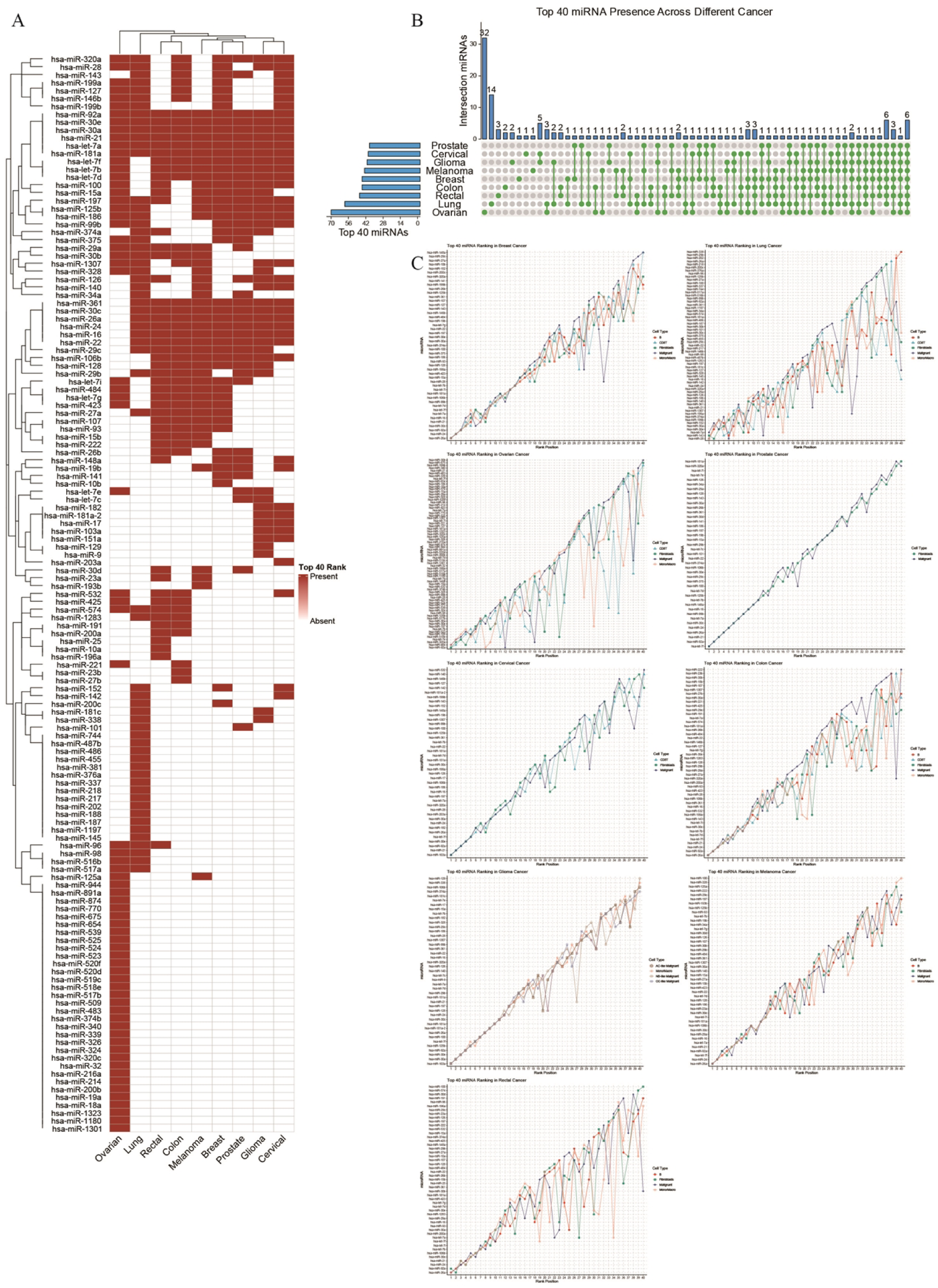
Pan-Cancer Spatial Comparison of miRNA Commonality and Heterogeneity. (A) Venn diagram illustrating six pan-cancer conserved miRNAs (e.g., hsa-miR-21, hsa-let-7a) consistently ranked in the top 40 across nine cancer types. (B) Heatmap displaying miRNA rankings across cancers. Red highlights conserved miRNAs (e.g., miR-30 family), while color gradients reflect cancer-specific variations. (C) Line plot showing average miRNA expression levels across cell types (fibroblasts, B cells, malignant cells) in nine cancers. Solid lines indicate conserved rankings in breast and cervical cancers; dashed lines highlight heterogeneity in lung and ovarian cancers.

This commonality highlights some shared molecular mechanisms across cancers, despite their distinct tissue origins and pathogenesis. The high expression of these miRNAs likely reflects the reliance of cancer cells on certain core signaling pathways. Subsequently, we computed the average expression of each miRNA across different cell types in the spatial transcriptomics (ST) data from 9 cancer types: breast cancer, cervical cancer, color cancer, glioma, lung cancer, melanoma, ovarian cancer, prostate cancer, and rectal cancer(**Figure 2C**). The miRNAs were ranked in ascending order based on their average expression levels, resulting in miRNA ranking lists for each cell type within each cancer. We then identified the top 20 miRNAs in each cancer type and cell type. Notably, the miRNA rankings exhibited a high degree of consistency across different cell types within each cancer. For instance, in breast cancer, the top three miRNAs—hsa-miR-26a, hsa-miR-24, and hsa-miR-92a—consistently ranked high in B cells, CD8T cells, fibroblasts, malignant cells, and monocytes/macrophages. In cervical cancer, hsa-miR-103a, hsa-miR-21, and hsa-miR-92a similarly ranked consistently across CD8T cells, fibroblasts, and malignant cells. In colon cancer, hsa-miR-26a, hsa-miR-92a, and hsa-miR-24 maintained a similar ranking across B cells, CD8T cells, fibroblasts, malignant cells, and monocytes/macrophages.

However, in lung cancer and ovarian cancer, the rankings of the top 20 miRNAs showed considerable inconsistency across different cell types. This lack of consistency suggests a high degree of heterogeneity in miRNA regulation within these cancer types, indicating that different cell types may exhibit distinct miRNA expression profiles. This variability could be influenced by several factors, including the molecular characteristics of the tumor, the complexity of the microenvironment, and the interactions between different cell types. The observed heterogeneity in lung and ovarian cancers may reflect the unique miRNA regulatory networks in these cancers, implying that miRNA functions and biological mechanisms may be more specific and complex within the different cellular contexts of these tumor types.

This analysis underscores both the shared and divergent roles of miRNAs across various cancer types. The identification of common miRNAs, such as hsa-miR-30a, hsa-miR-30e, hsa-miR-181a, hsa-let-7a, hsa-miR-92a, and hsa-miR-21, suggests that these miRNAs are involved in fundamental biological processes, such as cell proliferation, differentiation, and apoptosis, that transcend individual cancer types. The conserved role of these miRNAs across different cancers points to potential therapeutic targets for a broad range of malignancies.

Conversely, the observed heterogeneity in miRNA rankings, particularly in lung and ovarian cancers, highlights the complexity of miRNA regulation in the tumor microenvironment. This variability may be driven by the unique molecular signatures of each cancer type, as well as by the intricate interactions between different cellular components within the tumor. These findings suggest that miRNA-based therapies may need to be tailored to the specific cellular context and molecular features of each cancer type, particularly in tumors exhibiting significant heterogeneity.

In conclusion, the integration of miRNA expression data across multiple cancer types provides valuable insights into the common and distinct regulatory roles of miRNAs in cancer biology. These results underscore the importance of considering both the conserved and heterogeneous nature of miRNA networks when designing cancer therapeutics.

### 3. Pan-Cancer Spatial Analysis of miRNA Commonality in Cell Subtypes

To investigate the commonality of miRNA expression across cell subtypes in various cancer types, we analyzed fibroblasts, B cells, malignant cells, and monocytes/macrophages (Mono/Macro) in eight cancer types—Breast, Lung, Ovarian, Prostate, Cervical, Colon, Melanoma, and Rectal. Our analysis revealed several miRNAs consistently expressed across these cell subtypes, providing insights into their potential roles in cancer biology.

In fibroblasts, we identified several miRNAs highly expressed across all cancer types(**Figure 3A, B**). Notably, hsa-miR-30a and hsa-miR-30e were consistently ranked among the top 30 miRNAs in fibroblasts across all cancer types analyzed. Recent studies have shown that miR-30 family members can modulate fibroblast activation, which is crucial in tumor progression, especially in the context of fibrosis and the tumor microenvironment (TME). The expression of hsa-miR-30a and hsa-miR-30e in fibroblasts suggests they may play a pivotal role in regulating the fibrotic response in various cancers, impacting both tumor growth and metastasis(Eichelmann, Matuszcak, Hummel, & Haier, 2018; Wu et al., 2020).

**Figure 3.**
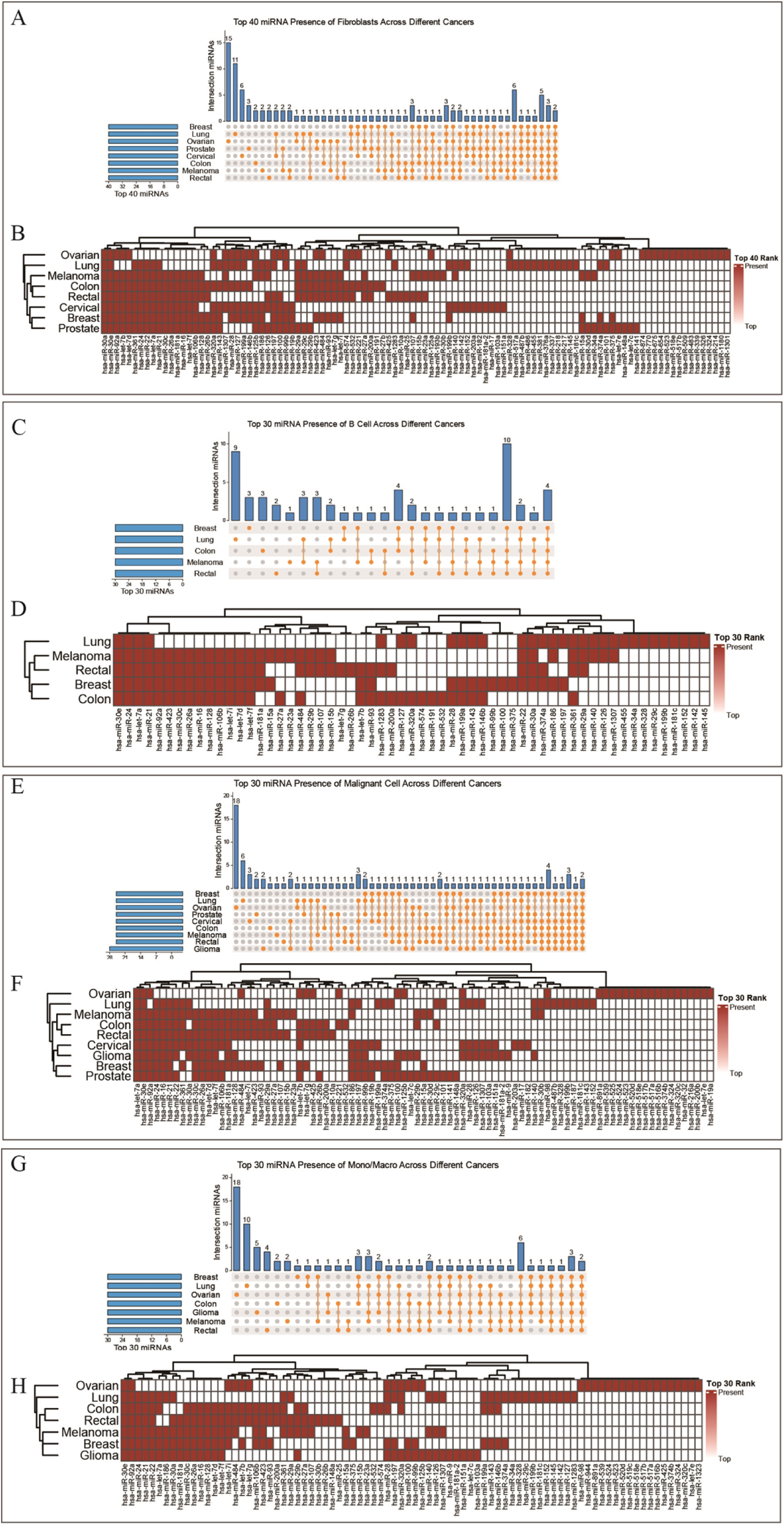
Pan-Cancer miRNA Conservation in Cell Subtypes. (A, B) Fibroblasts: Violin plots of hsa-miR-30a and hsa-miR-30e expression, conserved across cancers. These miRNAs regulate fibroblast activation and fibrosis. (C, D) B cells: Boxplots of hsa-let-7a (tumor suppressor) and hsa-miR-21 (oncogenic), showing balanced roles in immune modulation. (E, F) Malignant cells: Scatterplots of hsa-let-7a and hsa-miR-30e expression, linked to oncogene repression (e.g., RAS) and metastasis inhibition. (G, H) Monocytes/Macrophages: Heatmaps of miR-30e and miR-92a expression, indicating roles in macrophage polarization and immune response regulation.

In B cells, four miRNAs—hsa-let-7a, hsa-miR-21, hsa-miR-24, and hsa-miR-30e— were identified as consistently expressed across the five cancer types that included B cells (Breast, Lung, Colon, Melanoma, Rectal)(**Figure 3C, D**). The let-7 family, including hsa-let-7a, is a well-known tumor suppressor that regulates key oncogenes involved in cell proliferation and apoptosis(Eichelmann et al., 2018; Jamshidi et al., 2021). Meanwhile, hsa-miR-21 is an established oncogene, often upregulated in various cancers, where it promotes tumorigenesis by targeting anti-apoptotic pathways(Selcuklu, Donoghue, & Spillane, 2009). The consistent presence of hsa-miR-21, along with hsa-let-7a, highlights the balance between tumor suppression and promotion within the immune microenvironment, suggesting a complex interplay between B cells and the tumor microenvironment.

Among malignant cells, which were analyzed across nine cancer types, hsa-let-7a and hsa-miR-30e were the only miRNAs consistently found in the top 30(**Figure 3E, F**). These miRNAs’ roles in malignant cells are critical as hsa-let-7a is known to inhibit key oncogenes such as RAS(Lan et al., 2011), while hsa-miR-30e has been implicated in regulating cell migration and invasion(Yang et al., 2024). The consistent expression of these miRNAs in malignant cells across various cancer types underscores their potential as therapeutic targets, particularly in strategies aimed at modulating the tumor microenvironment and limiting metastasis.

Finally, in monocytes/macrophages (Mono/Macro), hsa-miR-30e and hsa-miR-92a were identified as the top common miRNAs across seven cancer types (Breast, Lung, Ovarian, Colon, Glioma, Melanoma, Rectal)(**Figure 3G, H**). MiR-30e has been reported to modulate the immune response, particularly in macrophages, by regulating the polarization of macrophages into either pro-inflammatory or anti-inflammatory phenotypes(Ahmad, Naqvi, Valverde, & Naqvi, 2023; Miranda et al., 2018). This suggests that miR-30e might play a critical role in shaping the immune landscape of tumors, potentially influencing the macrophage-mediated immune response to cancer cells(Conrad et al., 2023; X. Li, Guo, Min, Guo, & Zhang, 2019). On the other hand, miR-92a is associated with immune cell infiltration and tumor progression, with recent studies showing that it can influence macrophage activation and migration within the tumor microenvironment(Nilsson et al., 2012).

These findings reveal several common miRNAs that are consistently expressed across different cancer types and cell subtypes, providing important insights into the molecular mechanisms underlying tumor biology. The identification of hsa-miR-30a, hsa-miR-30e, and other key miRNAs in fibroblasts, B cells, malignant cells, and monocytes/macrophages suggests that these miRNAs may be involved in fundamental processes such as immune modulation, tumor progression, and the regulation of the tumor microenvironment. Furthermore, the consistent expression of miRNAs like hsa-let-7a and hsa-miR-21 across immune and malignant cell types reflects their dual role in both tumor suppression and promotion, underscoring the complex interplay of miRNAs in cancer.

Interestingly, the differences observed in miRNA expression across cancer types, such as the variation in Mono/Macro cell miRNAs between cancer types, further highlight the heterogeneity of miRNA regulation within the tumor microenvironment. These findings underscore the need to consider both common and unique miRNA signatures when designing therapeutic strategies targeting the tumor microenvironment.

In conclusion, this pan-cancer spatial analysis of miRNA expression across various cell subtypes reveals important insights into the shared and distinct roles of miRNAs in cancer. These results have significant implications for the development of miRNA-based therapeutics, particularly in tailoring treatments that modulate the tumor microenvironment or immune response.

### 4. Comparative Analysis of miRNA Signatures Across Cancer Types and Cell Subtypes

To explore the role of cell-type-specific miRNAs across different cancer types, we conducted a comprehensive analysis of miRNA expression profiles from nine cancer types, identifying top 50 differentially expressed miRNAs. Our findings reveal significant variations in miRNA signatures across various cell subtypes, highlighting the influence of miRNAs in cancer progression, immune regulation, and stromal interactions.

In breast cancer, we observed that hsa-miR-30a is specifically expressed in B cells(**Figure 4A**). Additionally, hsa-miR-146b, upregulated in B cells, CD8+ T cells, and monocytes/macrophages, plays a crucial role in inflammatory responses and immune regulation. Other miRNAs such as hsa-miR-22 (CD8+ T-cell-specific) and hsa-miR-199b (fibroblast-specific) further illustrate the complex interaction between immune and stromal components in the tumor microenvironment (TME). Malignant cells across multiple cancers shared several miRNAs, including hsa-miR-141, hsa-miR-29c, and hsa-miR-93, which are associated with cell proliferation and invasion, underscoring the universal role of these miRNAs in oncogenesis.

**Figure 4.**
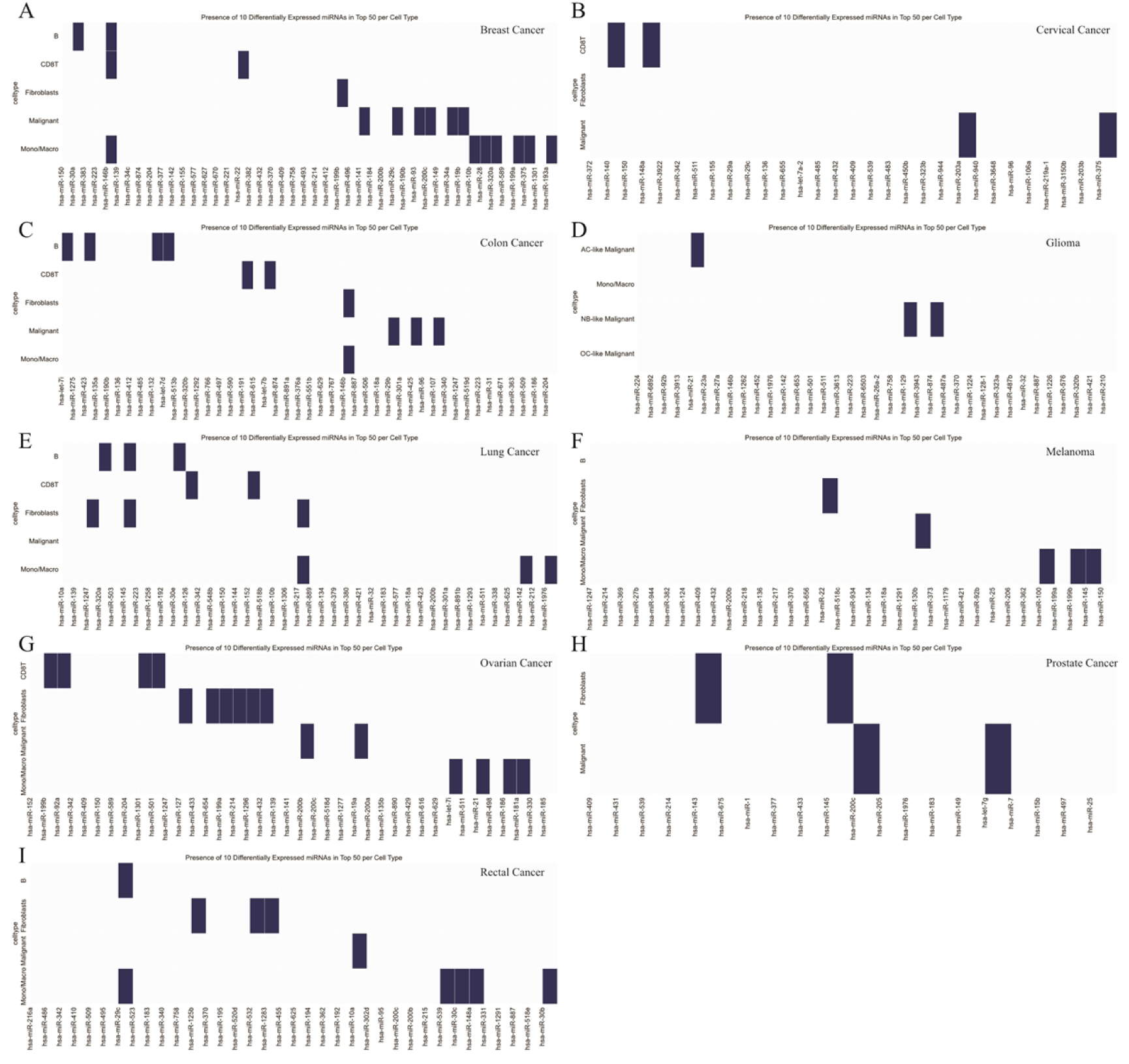
Cell-Type-Specific miRNA Signatures Across Cancers. **(A) Breast cancer**: hsa-miR-30a (B cells), hsa-miR-146b (B cells, CD8+ T cells, monocytes/macrophages), hsa-miR-22 (CD8+ T cells), and hsa-miR-199b (fibroblasts) delineate immune-stromal crosstalk. **(B) Cervical cancer**: CD8+ T cells express hsa-miR-140 and hsa-miR-148a (immune surveillance), while hsa-miR-203a (differentiation) and hsa-miR-375 (proliferation) dominate malignant cells. **(C) Colon cancer**: B cells exhibit hsa-let-7i, miR-423, and miR-132 (tumor suppression), with hsa-miR-146b shared across fibroblasts and monocytes/macrophages (inflammation/remodeling). **(D) Glioma**: Subtype-specific signatures include hsa-miR-21 (AC-like) and hsa-miR-129 (NB-like), indicating divergent oncogenic pathways. **(E) Lung cancer**: hsa-miR-320a/hsa-miR-145 (B cells), hsa-miR-217 (fibroblast activation), and hsa-miR-200c (malignant cells, EMT) highlight microenvironmental interplay. **(F) Melanoma**: hsa-miR-22 (fibroblasts) and hsa-miR-130b (malignant cells, metastasis) underscore stromal-tumor dynamics. **(G) Ovarian cancer**: Fibroblast-enriched miRNAs (hsa-miR-127, hsa-miR-654, hsa-miR-199a) and hsa-miR-21 (dual stromal/immune roles) reflect TME complexity. **(H) Prostate cancer**: hsa-miR-143/hsa-miR-145 mark fibroblasts, linked to desmoplastic stroma. **(I) Rectal cancer**: hsa-miR-125b/hsa-miR-532 (fibroblasts), hsa-miR-10a (malignant cells), and hsa-miR-539 (monocytes/macrophages) implicate inflammation-driven TME remodeling.

In cervical cancer, hsa-miR-140 and hsa-miR-148a were identified as CD8+ T-cell markers, suggesting their involvement in immune surveillance(**Figure 4B**). Moreover, hsa-miR-203a, associated with differentiation, and hsa-miR-375, linked to cell growth and survival, were highly expressed in malignant cells, reinforcing the contribution of miRNAs in tumor progression.

For colon cancer, distinct miRNA signatures were observed in B cells and CD8+ T cells(**Figure 4C**). Specifically, hsa-let-7i, miR-423, and miR-132 were identified as B-cell markers, with the let-7 family known for its tumor-suppressive effects. Interestingly, hsa-miR-146b was shared between fibroblasts and monocytes/macrophages, indicating its role in inflammation and tissue remodeling across different cancer types.

In glioma, miRNA profiles were subtype-specific, with hsa-miR-21 marking AC-like malignant cells and hsa-miR-129 linked to NB-like malignancies, suggesting distinct miRNA-driven pathways in glioma subtypes(**Figure 4D**).

In lung cancer, various miRNAs marked immune cells and fibroblasts, including hsa-miR-320a and hsa-miR-145 in B cells, and hsa-miR-1247 and hsa-miR-145 in fibroblasts, emphasizing the intricate interactions between immune and stromal cells in tumor progression(**Figure 4E**). Furthermore, hsa-miR-217, a marker of fibroblasts, is involved in fibroblast activation and ECM remodeling, while hsa-miR-200c marks malignant cells and regulates epithelial-mesenchymal transition (EMT), a key process in metastasis(Zhao Lin et al., 2024).

In melanoma, hsa-miR-22 and hsa-miR-130b were identified as fibroblast and malignant cell markers, respectively, with miR-130b known to promote metastasis through matrix metalloproteinase regulation(**Figure 4F**).

In ovarian cancer, a wide array of miRNAs, such as hsa-miR-127, hsa-miR-654, and hsa-miR-199a, marked fibroblasts, reinforcing the importance of the TME in tumor progression(**Figure 4G**). Notably, hsa-miR-21, present in both monocytes/macrophages and fibroblasts, suggests its role in regulating inflammation and stromal interactions.

In prostate cancer, hsa-miR-143 and hsa-miR-145 were key markers for fibroblasts, reflecting the desmoplastic reaction that is a hallmark of prostate cancer(**Figure 4H**). In rectal cancer, hsa-miR-125b and hsa-miR-532 marked fibroblasts, while hsa-miR-10a identified malignant cells(**Figure 4I**). Additionally, hsa-miR-539, a marker for monocytes/macrophages, further emphasized the role of inflammation in the TME. These results reveal a complex and cancer-specific distribution of miRNAs within different cell types of the TME, highlighting the diverse roles miRNAs play in regulating tumor growth, immune evasion, and stromal remodeling. Further studies into these miRNA signatures may pave the way for more targeted and precise therapeutic interventions in cancer.

### 5. Validation of miRNA - related approach in breast cancer case study and analysis of their regulatory roles in different cell types

To validate the effectiveness of our approach, we conducted a case study on breast cancer, focusing on the top ten miRNAs that were significantly differentially expressed in four specific cell types in the tumor microenvironment: malignant cells, B cells, fibroblasts, and myofibroblasts. We selected the hallmark miRNAs for malignant cells, B cells, fibroblasts, and myofibroblasts based on the criteria of P.adj<0.05 and the top 10 LogFC ranking. We found that the miRNAs enriched in malignant cells, B cells, fibroblasts, and myofibroblasts are all significantly associated with breast cancer (**Figure 5A**). This indicates that the enriched miRNAs possess disease specificity. Additionally, we predicted the target genes and miRNA-target gene interaction patterns for these hallmark miRNAs in the four cell types (**Figure 5B**).Within the malignant cells, our analysis reveals a positive regulatory relationship between miR-205 and the androgen receptor (AR) gene. This finding aligns with the study by (Kalinina et al., 2020), who proposed that the expression levels of AR and miR-205 could serve as predictive markers for lymph node metastasis in specific breast cancer subtypes. Furthermore, (Yu et al., 2013) demonstrated an inverse correlation between YAP1 and miR-200a expression in breast cancer clinical specimens, with miR-200a expression associated with distant metastasis in breast cancer patients. Additionally, (Sun et al., 2014) reported that miR-200b directly inhibits the expression of lysyl oxidase (LOX), consequently reducing tumor invasion. These findings collectively underscore the complex and diverse roles of miRNAs in breast cancer progression and metastasis. In B cells, (Cui et al., 2022) found that targeted OMV(tRNA-pre-miR-126) treatment inhibited the expression of oncogenic CXCR4 and significantly restrained breast cancer tissue proliferation. (Zhu et al., 2011) reported that miR-126 targets both VEGFA and PIK3R2, and its downregulation in tumors may increase VEGF/PI3K/AKT signaling pathway activity, inducing vascular formation, tumor growth, and metastasis. In fibroblasts, (Wu et al., 2018) demonstrated that miR-199b-5p inhibits triple-negative breast cancer cell proliferation, migration, and invasion by targeting DDR1. Specifically, miR-199b-5p inhibited DDR1 gene expression by directly targeting its 3′ untranslated region (UTR).

**Figure 5.**
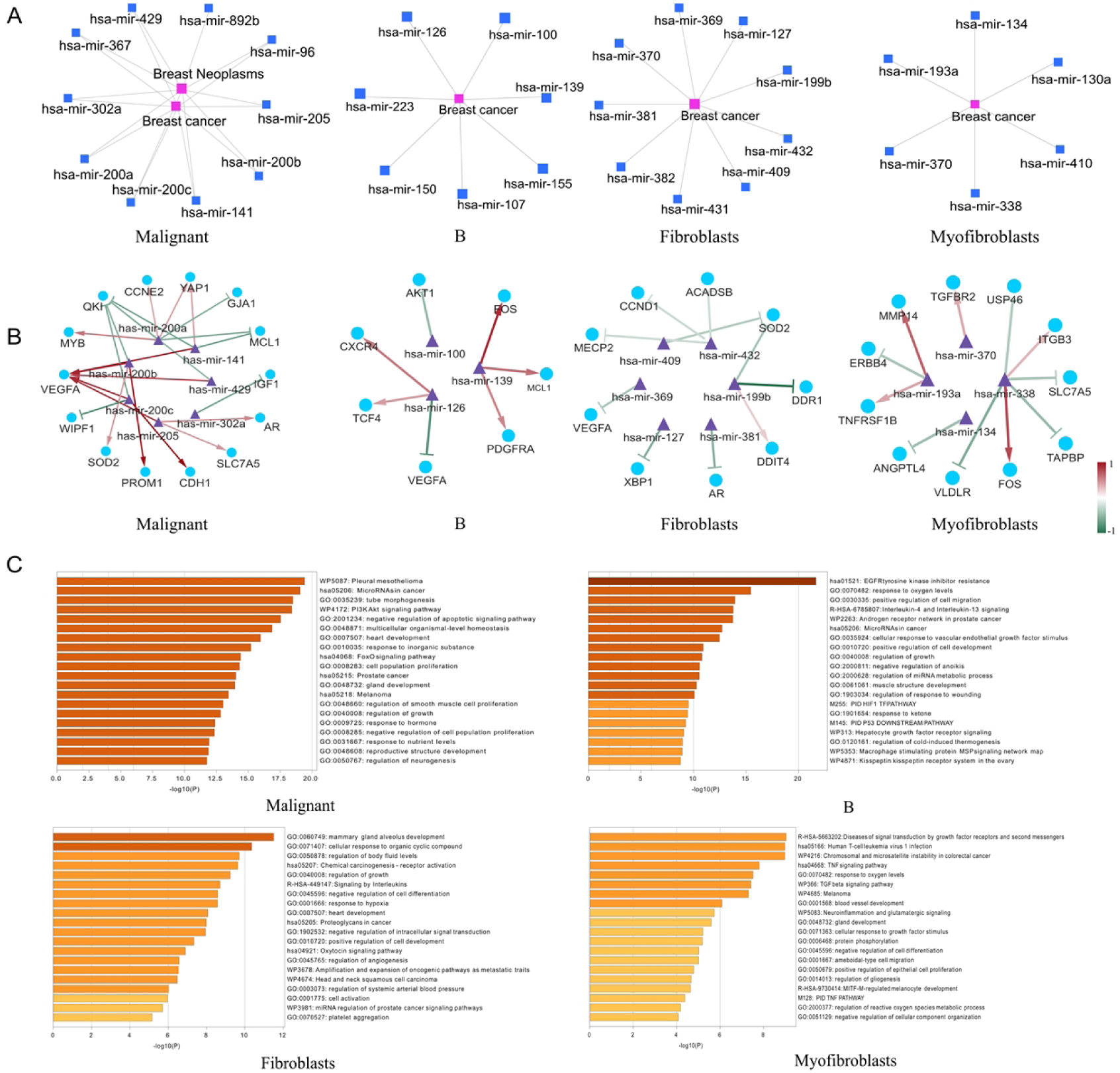
Breast cancer top miRNAs-target gene and miRNAs-disease regulatory network. (A) Network diagram linking top differentially expressed miRNAs (malignant cells, B cells, fibroblasts, myofibroblasts) to breast cancer pathways. (B) miRNA-target interaction map: Red edges (positive Pearson correlation, e.g., miR-205-AR), green edges (negative correlation, e.g., miR-200b-LOX). Line darkness reflects correlation strength. (C) Pathway enrichment bar charts: Malignant cells enrich “PI3K-Akt signaling,” B cells associate with “EGFR resistance,” fibroblasts link to “mammary gland development,” and myofibroblasts to “TNF signaling.”

To elucidate the regulatory relationships between miRNAs and their target genes, we performed a Pearson correlation analysis based on their expression levels. Our results reveal distinct regulatory patterns in various cell types, showing both upregulation and downregulation of miRNAs relative to their target genes in breast neoplasms (**Figure 5C**). In malignant cells, the target genes of enriched miRNAs are primarily involved in pathways such as “MicroRNAs in cancer,” “tube morphogenesis,” and the “PI3K Akt signaling pathway.” In B cells, the target genes are mainly associated with “EGFR tyrosine kinase inhibitor resistance” and “response to oxygen levels.” For fibroblasts, the enriched miRNA target genes are involved in “mammary gland alveolus development” and “cellular response to organic cyclic compounds.” In myofibroblasts, the target genes are predominantly linked to “Diseases of signal transduction by growth factor receptors and second messengers,” “Human T-cell leukemia virus 1 infection,” and the “TNF signaling pathway.” We observed that many of these genes are associated with miRNA regulation in cancer. This may be due to the presence of malignant cells mixed within each spot during our deconvolution process. miRNAs serve as mediators of intercellular signaling between malignant and other cell types, suggesting that the miRNAs assessed in spatial transcriptomics are more likely to represent intercellular interactions rather than cell type-specific miRNA characteristics. These findings are consistent with previous studies and provide further insight into the complex regulatory mechanisms of miRNAs in breast cancer.

## Discussion

In this study, we developed STmiR, a tool designed to predict miRNA activity from spatial transcriptomics (ST) data, overcoming some of the major challenges associated with miRNA analysis in spatial contexts. Through the application of this tool, we constructed a predictive model based on bulk RNA-seq data, specifically using paired miRNA-mRNA expression profiles from four major cancer types—Breast Cancer, Non-Small Cell Lung Cancer, Ovarian Cancer, and Prostate Adenocarcinoma— available in The Cancer Genome Atlas (TCGA). By leveraging the power of machine learning, particularly the XGBoost model, we demonstrated that our tool could predict miRNA expression with a high degree of accuracy, achieving a correlation greater than 0.8 between predicted and actual miRNA levels. This performance confirms the potential of STmiR for studying miRNA spatial distribution and gene regulation in cancer.

Our work highlights both the promise and challenges of using spatial transcriptomics to study miRNA regulation. While miRNAs are crucial regulators in gene expression and cellular communication within the tumor microenvironment, their small size and low abundance present significant technical barriers to their analysis in spatial transcriptomics. Despite these challenges, increasing evidence from recent studies underscores the importance of miRNAs in regulating key processes such as tumor progression, immune modulation, and metastasis. The spatial distribution of miRNAs, which was previously difficult to study, is now more accessible due to the development of tools like STmiR.

The predictive model in STmiR capitalizes on the understanding that gene expression data from different modalities are often interrelated. Previous studies have shown that gene expression can serve as a powerful predictor for other molecular features, such as transcription factor activity and copy number variations, at the single-cell level. This notion is central to STmiR, as it uses mRNA expression data to predict miRNA activity. Given that miRNAs regulate their target genes through specific sequence binding, this approach is both feasible and effective. Furthermore, our results show that the miRNAs predicted by STmiR correlate with biologically relevant processes in specific cancer types. For example, high expression of miR-200b in malignant breast cancer cells was linked to epithelial-mesenchymal transition, lymph node metastasis, and chemotherapy resistance, all of which are critical in cancer progression.

Despite the strengths of STmiR, our study also presents several limitations that must be addressed in future work. One of the main limitations of this research is the sparsity of spatial transcriptomic data. Spatial transcriptomics, while a promising technology, has limited resolution in terms of the number of regions sampled, which can result in data that is less comprehensive than bulk RNA-seq. This limitation makes it more difficult to obtain accurate spatially resolved predictions of miRNA activity across tissue structures. Additionally, the model’s training on bulk RNA-seq data might introduce biases due to the inherent differences between bulk tissue samples and the high-resolution, spatially defined data typical of ST analyses. To overcome these limitations, future iterations of STmiR should incorporate more extensive datasets, including larger cohorts and spatially resolved data from diverse cancer types, to improve its generalizability and accuracy.

Another challenge lies in the complexity of the tumor microenvironment, which varies widely across different cancer types and even within regions of the same tumor. The heterogeneity observed in the spatial distribution of miRNAs, as revealed by our study, suggests that miRNA functions may be highly context-dependent, varying not only between tumor types but also across cell types within the same tumor. For instance, miRNAs such as hsa-miR-30a and hsa-miR-21 exhibited distinct spatial patterns across fibroblasts, B cells, and malignant cells, indicating their specialized roles in different cellular contexts. These findings emphasize the need to refine our understanding of miRNA functions, considering the spatial and cellular heterogeneity in tumors.

Despite these challenges, our research offers important contributions to the field of spatial miRNA analysis. By providing a novel tool for predicting miRNA activity in the spatial context, STmiR opens new avenues for studying the regulatory roles of miRNAs in cancer biology. The ability to investigate miRNA expression at both the cellular and spatial levels will be invaluable in uncovering the mechanisms underlying tumorigenesis, metastasis, and response to treatment. Moreover, the insights gained from this tool could guide the development of new therapeutic strategies that target miRNA-mediated pathways, offering hope for more effective and personalized cancer treatments.

In conclusion, STmiR represents a significant step forward in understanding the spatial regulation of miRNAs in cancer. The tool’s ability to predict miRNA activity in a spatial context provides a powerful resource for exploring the intricate networks that govern gene expression in complex tissues. While there are limitations to be addressed, the work presented here lays the foundation for future research into spatial miRNA dynamics and their roles in cancer progression and therapy.

## Resources and Methods

### Key Resources table

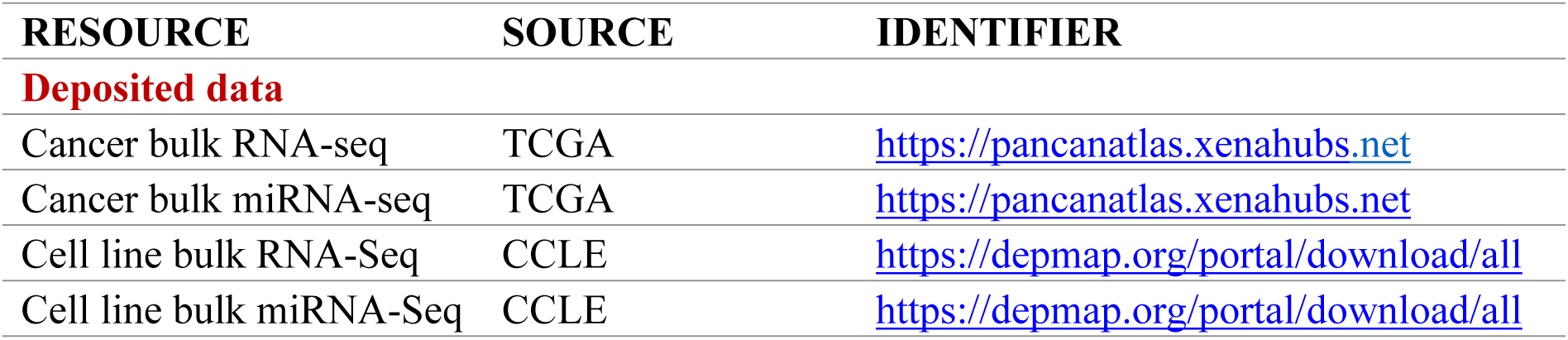
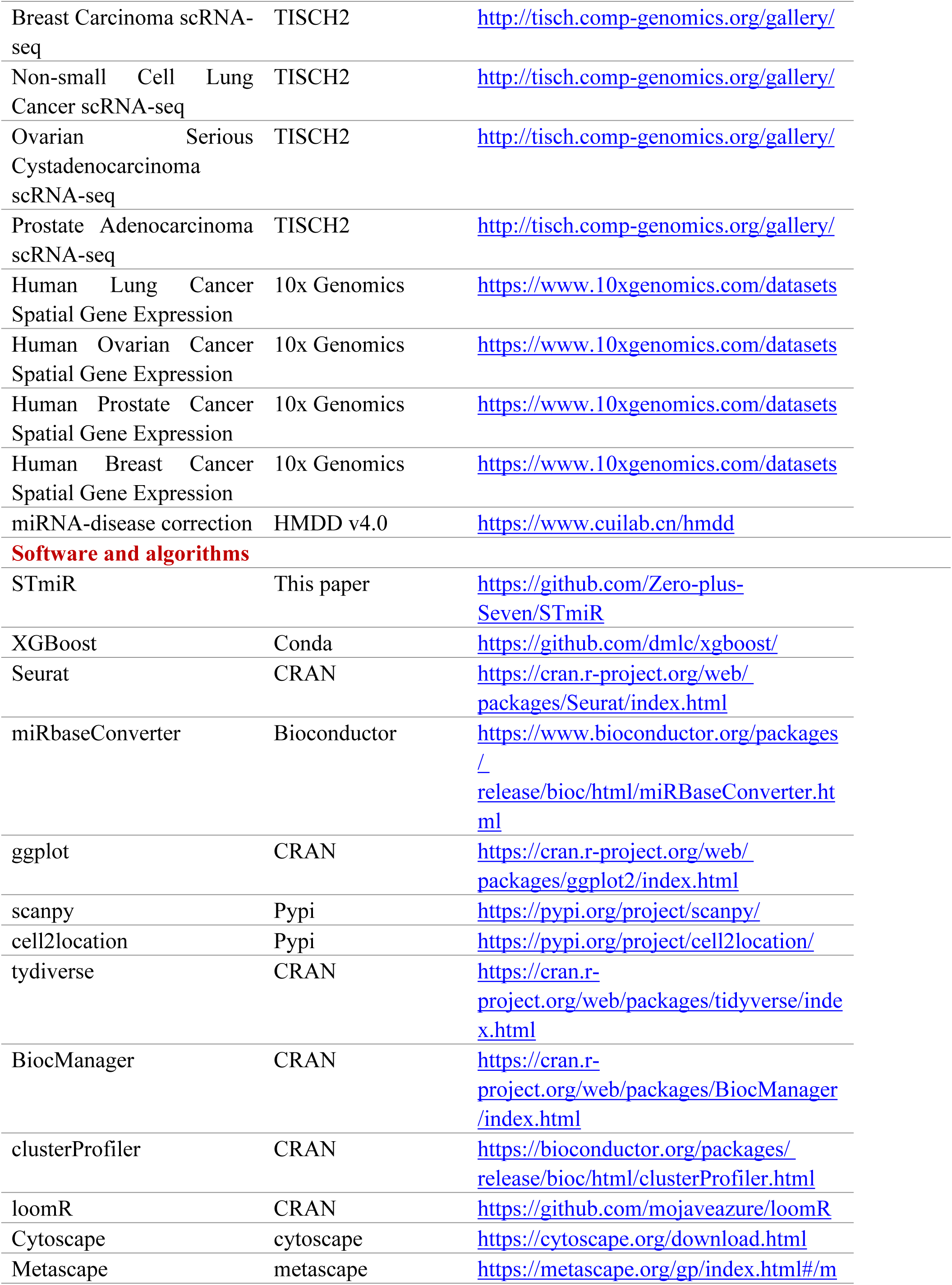
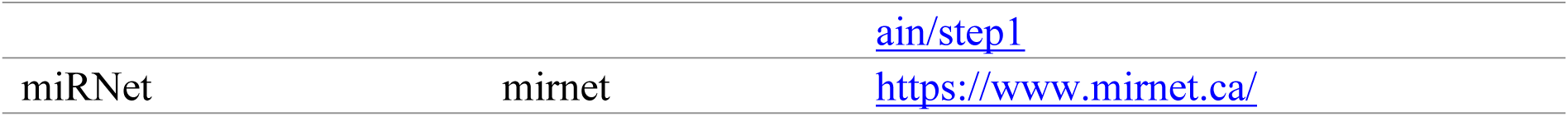

## Declaration

### Ethics approval and consent to participate

Not applicable.

### Consent for publication

Not applicable.

### Availability of data and materials

This paper analyzes existing, publicly available data. These accession numbers for the datasets are listed in the Key Resources table. Additionally, the original code used in this study was publicly deposited on GitHub, and its link are listed in the Key Resources table.

### Competing interests

The authors declare no competing interests.

### Funding

This work was supported in part by the National Natural Science Foundation of China (No. 62072128, 62002079 and 62202112), the Natural Science Foundation of Guangdong Province of China (No. 2023A1515011401), the National key R and D Program of China(Grant 2019YFA0706338402), the Municipal School Joint Fund of Guangzhou Science and Technology Bureau (SL2022A03J00935), and the Open Project of Guangdong Provincial Key Laboratory of Artificial Intelligence in Medical Image Analysis and Application (No. 2022B1212010011).

### Authors’ contributions

Jiaqi Yuan: Data curation, Conceptualization, Methodology, Software Writing-Original draft preparation. Peng Xu: Formal analysis, Supervision, Project administration, Funding acquisition. Wenbin Liu: Software, Validation, Writing - Review & Editing, Funding acquisition. Zheng Ye: Investigation, Visualization, Writing-Reviewing and Editing, Supervision.

## Acknowledgements

Not applicable.

## Notes

### Competing Interest Statement

The authors have declared no competing interest.

